# LigRNA: High-accuracy prediction of RNA-ligand binding sites via dynamic chemical complementarity and language model encoding

**DOI:** 10.64898/2026.10.15.759315

**Authors:** Xin Wang, Heqin Zhu, Hao Li, Ao Chang, Feng Zhang, Haobin Chen, Chenxi Wang, Ke Chen, Shaohua Kevin Zhou, Peng Xiong

**Author notes:** Corresponding authors: Peng Xiong, Shaohua Kevin Zhou, Heqin Zhu. Xin Wang and Heqin Zhu contributed equally to this work.

## Abstract

RNA-binding small molecules represent a validated therapeutic modality, yet pinpointing their exact RNA binding sites remains critical for mechanistic understanding and drug optimization. Experimental methods, while precise, are costly and low-throughput, necessitating robust computational alternatives. Approaches based on static feature concatenation may overlook input-dependent relationships between RNA residues and ligand functional groups. Here we present LigRNA, a graph neural network framework that predicts RNA-ligand binding sites through joint cross-modal encoding of both molecules. LigRNA integrates structure-guided language model embeddings with physicochemical features-solvent-accessible surface area and secondary structure types, to construct comprehensive RNA representations. Ligands are encoded as attributed molecular graphs via graph attention networks, preserving atom-level chemical topology rather than compressing molecules into fixed fingerprints. A cross-attention mechanism then enables dynamic, ligand-dependent modulation of RNA nucleotide attention, capturing context-specific interaction patterns. Benchmarked on TE9 and TE18 datasets, LigRNA achieves F1 scores of 0.73 and 0.58, improving on GATRsite by 7.4% and 18.4%, respectively. Ablation studies identify the RNA residual encoder as the most consistent contributor across the two benchmarks. The source code is freely available at https://github.com/WillowWang0216/LigRNA.

## 1. Introduction

Ribonucleic acid (RNA) has emerged as a compelling class of therapeutic targets for small-molecule drugs, with recent clinical successes validating that structured RNA elements can be drugged by drug-like small molecules (Chen et al., 2024; Warner et al., 2018). These compounds exert their therapeutic effects by engaging specific regions of functional RNA structures; accordingly, accurate identification of RNA-ligand binding sites is a prerequisite for lead optimization and mechanistic understanding in RNA-targeted drug discovery (Disney et al., 2016; Warner et al., 2018). While experimental techniques such as X-ray crystallography and cryo-electron microscopy can determine binding sites with atomic resolution, their high cost and limited throughput preclude systematic screening across large chemical libraries, creating a pressing need for accurate computational prediction methods.

Computational prediction of RNA-ligand binding sites has advanced considerably over the past decade. Early geometry-based methods such as Rsite (Zeng et al., 2015) and Rsite2 (Zeng and Cui, 2016) localize candidate functional sites using distances between RNA nucleotides derived from tertiary and secondary structures, respectively. The network-based approach RBind (Wang et al., 2018) uses nucleotide interaction networks to identify binding sites, whereas RNAsite (Su et al., 2021) combines sequence-profile and structural features with random forest classifiers. While these methods demonstrated the feasibility of computational prediction, their reliance on hand-crafted rules or shallow classifiers may limit generalization to novel RNA-ligand pairs. Deep learning has since reshaped the landscape. RLBind (Wang et al., 2023) showed that jointly processing full-length RNA sequences and local nucleotide neighborhoods improves performance, and GATRsite (Sun et al., 2025) further extended this direction by integrating pre-trained RNA language model embeddings with graph attention networks, achieving state-of-the-art results on the TE9 and TE18 benchmarks. Geometric deep learning methods incorporating RNA language models have also been explored for this task (Zhu et al., 2025c). Beyond binding site prediction, related computational methods for RNA-small molecule binding affinity prediction have also been developed; for instance, RLaffinity integrates RNA pocket and small molecule information through a 3D convolutional neural network with contrastive pre-training (Sun and Gao, 2024). Nevertheless, two persistent bottlenecks continue to constrain prediction accuracy.

First, encoding a ligand as a single fixed-length vector limits access to individual atom representations during RNA-ligand feature fusion. This can obscure atom-level chemical information of the ligand—information that fundamentally governs hydrogen-bonding geometry, aromatic stacking, and electrostatic complementarity at the binding interface (Warner et al., 2018). Second, static concatenation or averaging does not explicitly provide nucleotide-specific weights over ligand atoms. Such fusion may be less suited to modeling the fact that binding modes are inherently ligand-dependent: different binding sites operate through distinct sets of hydrogen-bonding, aromatic-stacking, and electrostatic interactions, and a single modality may not adequately represent this diversity. Together, these considerations motivate atom-level ligand encoding and ligand-conditioned nucleotide representations.

To address these challenges, we present LigRNA, a graph neural network framework that predicts RNA-ligand binding sites from RNA sequences and small-molecule structures. On the ligand side, LigRNA represents each molecule as an attributed molecular graph processed through a graph attention network (Brody et al., 2022), preserving the full chemical topology of every atom rather than reducing the ligand to a fixed fingerprint—this enables the model to learn atom-type-specific matching patterns against RNA pockets using separate atom representations. On the RNA side, a residual convolutional network aggregates heterogeneous features—structure-guided language model embeddings (Zhu et al., 2025a), solvent-accessible surface area, and secondary structure types (Danaee et al., 2018)—over multi-nucleotide neighborhoods, allowing the model to perceive binding environments spanning typical pocket boundaries. Cross-attention then replaces static feature concatenation, enabling RNA nucleotides to update their representations conditioned on ligand atom features. Crucially, the entire pipeline operates on RNA sequences and secondary-structure annotations (Zhu et al., 2025b), enabling binding site prediction at early stages of drug discovery when tertiary structures may not yet be available.

## 2. Materials and Methods

This section describes the computational framework of LigRNA. We first detail the construction of training and test datasets from public RNA-ligand complex repositories. We then present the overall model architecture (Section 2.2), followed by detailed descriptions of each network component. Specifically, we describe the RNA feature extraction pipeline, the ligand feature extraction pipeline, the cross-attention interaction module, the training objectives and aggregation strategy, and finally the evaluation metrics.

### 2.1. Dataset Construction

The benchmark datasets used for training and evaluation are derived from two public repositories of RNA-ligand complex structures. The primary source is the Protein Data Bank (PDB) (Burley et al., 2021), from which we retain 78 RNA-ligand complexes following the PDB identifier list curated by RNAsite (Su et al., 2021). Ligands are defined as low-molecular-weight molecules (mass ≤ 1000 Da), encompassing metal ions, natural metabolites, and drug-like compounds; crystalline water molecules are explicitly excluded. For complexes harboring multiple ligands, each ligand’s binding site is preserved independently. Data cleaning follows the protocols established by RNAsite (Su et al., 2021) and RLBind (Wang et al., 2023), and RNA clustering is performed at a TM-score (Zhang and Skolnick, 2005; Altschul et al., 1990; Li and Godzik, 2006) threshold of 0.3 to ensure sequence diversity. This yields a training/validation set of 42 clusters (60 RNAs, Train60) and a held-out test set of 15 clusters (18 RNAs, TE18). A second test set (TE9) is derived from the RBind repository: after removing multi-chain or pseudo-inactive RNAs, 19 complexes remain, of which 10 overlap with Train60, leaving 9 non-redundant test cases.

To enrich the training corpus, an additional 228 RNA-ligand complexes were retrieved from the PDB as of October 2025, filtered by the same criteria as RNAsite and RLBind. These form the Train228 set (241 chains, 67 clusters), which shares no cluster overlap with either TE18 or TE9, eliminating data leakage. Train228 was merged with Train60 and split 8:2 into training and validation partitions. Although some RNA structures in Train228 are sequence-similar to Train60, the diversity of bound ligands introduces distinct micro-chemical interaction environments, which mitigate the class imbalance and data sparsity inherent to binding site prediction. Specifically, the training set contains 1,926 binding-site nucleotides and 5,703 non-binding nucleotides, while the test sets (TE9 and TE18) contain 328 binding-site nucleotides and 892 non-binding nucleotides. Full dataset composition and detailed statistics are provided in Supplementary Table S1, and the contained PDB structure identifiers are listed in Supplementary Table S2.

### 2.2. Model Architecture

The overall architecture of LigRNA is illustrated in Figure 1. The model takes an RNA sequence and a small-molecule ligand as input and outputs a binding-site probability for each nucleotide. It consists of four components: (A) an RNA encoder integrates structure-guided language model embeddings with explicit physicochemical features to produce a contextualized nucleotide representation; (B) a ligand encoder represents the molecule as an attributed molecular graph and extracts atom-level pharmacophoric features via graph attention; (C) a 3-layer multi-head cross-attention module updates RNA nucleotide representations using ligand atom features; (D) an MLP prediction head produces nucleotide probabilities through a sigmoid function; probabilities from two models are aggregated before thresholding. Each component is described in detail in the following subsections.

**Fig. 1:**
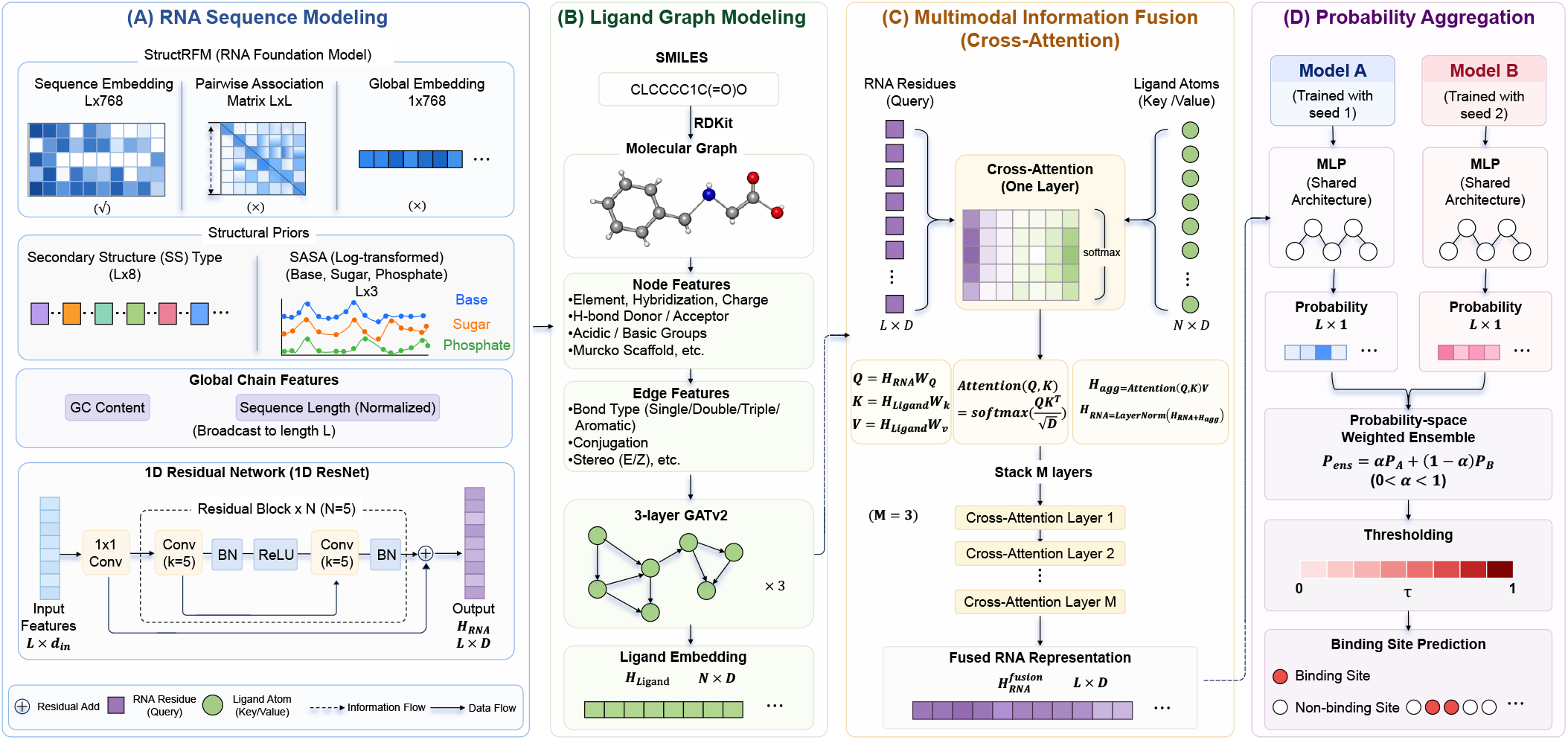
Schematic architecture of LigRNA. **(A) RNA Sequence Modeling.** The RNA branch takes as input the embedding features from the structRFM model, augmented with structural prior knowledge including secondary structure (SS) types and SASA. These features are processed through a 1D Residual Network (1D ResNet) with residual blocks to capture local sequence contexts. **(B) Ligand Graph Modeling**. Ligands are represented as attributed molecular graphs derived from SMILES strings. A three-layer GATv2 graph neural network is employed to extract atom-level topological and physicochemical features, producing enhanced ligand embedding representations (*H*_Ligand_). **(C) Multimodal Information Fusion**. A 3-layer multi-head cross-attention mechanism facilitates dynamic interaction between the two modalities, where RNA residues serve as the Query and ligand atoms act as the Key/Value. This process conditions nucleotide representations on ligand atom features. **(D) Probability Aggregation**. Each model (Model A and Model B) produces nucleotide probabilities through a Multi-Layer Perceptron (MLP) and sigmoid function. The two probability outputs are combined by a weighted average and thresholded to identify binding and non-binding sites.

### 2.3. RNA Feature Representations

As illustrated in Figure 1A, the RNA branch takes as input the embedding features from the structRFM model, augmented with structural prior knowledge including secondary structure types and SASA. RNA binding-site patterns are associated with evolutionary sequence variation and base-pairing patterns: nucleotides involved in ligand recognition are subject to distinct selective pressures and frequently exhibit non-canonical pairing geometries that distinguish them from structurally inert positions. Capturing these signals from the sequence alone, however, is challenging without a large-scale evolutionary context. To this end, LigRNA leverages structRFM (Zhu et al., 2025a), a structure-guided RNA foundation model pre-trained on approximately 21 million RNA sequences and structures via structure-guided masked language modeling (SgMLM). Unlike standard sequence-only language models, structRFM learns to co-predict sequence tokens and base-pairing partners in a joint unsupervised objective, thereby internalizing deep evolutionary co-variation and long-range structural dependencies beyond the information available at individual positions. The resulting embeddings may be informative for binding site prediction: nucleotides at functional interfaces tend to exhibit distinctive co-variation signatures and contextual structural preferences that structRFM may capture even in the absence of experimental structural data.

Specifically, structRFM generates three complementary outputs. The nucleotide-level embeddings (*L ×* 768) encode rich contextual sequence semantics and latent structural preferences, serving as the primary feature input for our downstream residual network. The pairwise structural association matrix (*L × L*) provides base-pairing probabilities inferred directly from sequence during pretraining. The globally pooled embedding (1 *×* 768) is a single chain-level vector designed for sequence-level tasks and does not carry sufficient granularity for fine-grained per-nucleotide binding site prediction. Consequently, only the nucleotide-level embeddings from structRFM are incorporated into LigRNA.

#### 2.3.1. Physicochemical Structure Features

Pre-trained RNA language models excel at capturing evolutionary sequence patterns but offer limited explicit representation of the three-dimensional physicochemical environment that directly governs molecular recognition. To address this, LigRNA extracts three categories of engineered physicochemical features grounded in established molecular recognition principles: SASA, secondary structure types, and global chain properties (Warner et al., 2018).

#### 2.3.2. Solvent-Accessible Surface Area (SASA)

The SASA of a nucleotide directly reflects its exposure to the solvent and therefore its potential to interact with incoming ligand molecules. Nucleotides buried in the RNA core, protected by the dense sugar-phosphate backbone, may be less accessible to incoming ligands, whereas surface-exposed nucleotides in grooves and pockets form the functional surface of the binding interface. To operationalize this principle, the SASA of each nucleotide was computed with PyMOL and further partitioned into three chemical subregions—base, sugar ring, and phosphate group— reflecting the distinct physicochemical character of each surface component. Partitioning distinguishes these components because the base face, sugar edge, and phosphate group contribute differentially to *π*-*π* stacking, hydrogen bonding, and electrostatic interactions with ligand atoms (Morgan et al., 2017). A logarithmic transformation was applied to the raw area values to suppress the influence of extreme outliers, yielding a feature tensor of dimension *L ×* 3.

#### 2.3.3. Secondary Structure Types

The types of RNA secondary structure: internal loops, bulges, and hairpin loops are enriched in unpaired nucleotides that present Lewis-basic or acidic groups capable of hydrogen bonding, whereas ligand accessibility in helical regions depends on groove geometry. To encode these topological propensities, each nucleotide was classified into one of eight secondary structure element types (paired helix, internal loop, bulge, hairpin loop, multi-branch junction, knotting/pseudoknot, chain break, or other) by parsing the dot-bracket notation (DBN) of the RNA structure (Danaee et al., 2018). Spatial consistency of the parsed secondary structure was verified against the experimental coordinates using PyMOL, and one-hot encoding produces an 8-dimensional topological vector per nucleotide. Notably, an analysis of binding site composition in the TE9 dataset (Fig. 2) describes the proportions of different motifs: knotted and pseudoknot structures account for 37.3%, internal loops for 18.1%, and unclassified regions for 23.2%. In contrast, conventional secondary structure elements (hairpin loops 8.1%, helices 7.2%, bulges 5.7%) account for smaller fractions of the annotated sites. These percentages describe binding-site composition; establishing motif enrichment requires comparison with motif frequencies among all nucleotides.

**Fig. 2:**
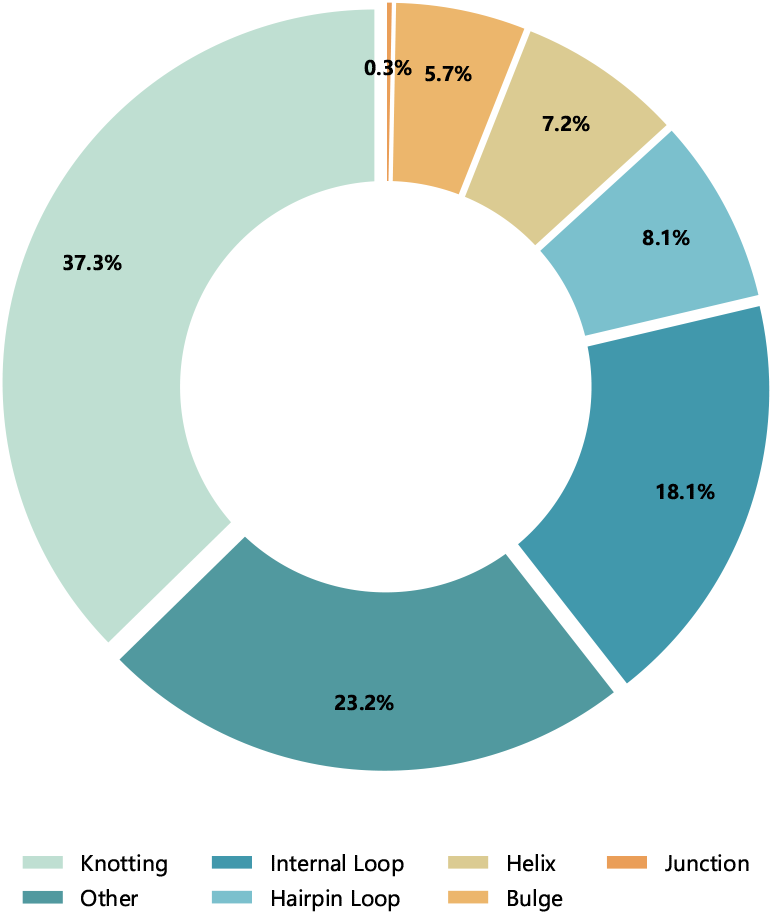
Distribution of RNA secondary structure types among binding-site nucleotides in TE9. The pie chart illustrates the proportion of each secondary structure motif among binding site nucleotides in the TE9 dataset. Knotting/pseudoknot annotations (37.3%) constitute the largest category, followed by unclassified motifs (23.2%) and internal loops (18.1%). Hairpin loops, helices, bulges and multi-branch loops together account for approximately 21.3%.

#### 2.3.4. Global Chain Features

Beyond local physicochemical properties, the thermodynamic stability and overall folding landscape of the entire RNA molecule provide important contextual priors for binding site prediction. Two global chain-level descriptors are computed and broadcast as scalar features to each individual nucleotide position. The first descriptor, mean GC content, serves as a coarse proxy for sequence-dependent thermodynamic stability: G-C base pairs form three hydrogen bonds compared to two for A-U pairs, contributing to higher melting temperatures in GC-rich regions (Takahashi and Sugimoto, 2020). However, the relationship between GC content and global structural compactness is context-dependent and modulated by ionic conditions and non-canonical interactions. Mean GC content is computed as the proportion of G and C nucleotides across the full sequence and broadcast uniformly to all positions. The second descriptor, normalized sequence length, characterizes overall structural complexity and conformational entropy: longer RNAs possess more degrees of freedom during folding and are statistically more likely to form intricate tertiary structures with buried binding pockets, whereas shorter RNAs tend to exhibit proportionally larger surface-to-volume ratios and simpler topologies (Takahashi and Sugimoto, 2020). Normalized sequence length is computed as the raw length divided by the maximum sequence length in the training corpus, and broadcast as a constant scalar to all positions.

#### 2.3.5. RNA Encoder

The RNA feature vector (structRFM embeddings concatenated with SASA, secondary structure, and global chain features) is first projected to a common hidden dimension *D* via a 1 *×* 1 convolutional layer. Five residual blocks are then stacked, each containing two convolutional layers with batch normalization (BN), ReLU activation, and a dropout layer. Each block performs:

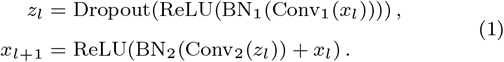

where Conv_1_ and Conv_2_ are both 1D convolutions with kernel size *k* = 5, with stride 1, dilation 1, and length-preserving padding. Across five blocks (ten convolutions), the nominal receptive field is 1 + 10(5 ™ 1) = 41 positions in the input feature sequence. The structRFM embeddings may already contain longer-range information. The skip connection (+ *x*_*l*_) mitigates gradient degradation in deep networks.

### 2.4. Ligand Feature Representations

As illustrated in Figure 1B, the ligand branch represents each molecule as an attributed molecular graph and processes it through a three-layer GATv2 graph neural network to extract atom-level topological and physicochemical features.

#### 2.4.1. Attributed Molecular Graph Construction

Ligand features were extracted using the open-source cheminformatics toolkit RDKit (RDKit contributors, n.d.). Each molecule was parsed from its SMILES string into an attributed molecular graph, where atoms are nodes and chemical bonds are edges. Atomic node features comprise three categories: intrinsic properties (element type, hybridization state, and formal charge); pharmacophoric attributes (hydrogen-bond donor/acceptor, acidic/basic character); and scaffold-based ring topology assignments (Table S3). Bond features include bond type (single, double, triple, aromatic), conjugation status, stereochemistry (*E/Z* configuration), and the bonded element pair (Table S4). All categorical features were discretized into index vectors and converted to PyTorch (Paszke et al., 2019) tensors. For monoatomic ions lacking covalent bonds (e.g., Mg^2+^), a dedicated preprocessing branch ensures numerical stability through placeholder graph construction.

#### 2.4.2. Atom-Level Ligand Encoder

Fixed-length fingerprints summarize ligands in a single vector without exposing separate atom representations to cross-attention. To capture atom-level chemical information, each ligand is represented as an attributed molecular graph and processed by a three-layer graph attention encoder (Brody et al., 2022), allowing each atom to attend differentially to its neighbors based on learned chemical compatibility. Critical pharmacophoric atoms—such as hydrogen-bond acceptors in heteroaromatic rings or charged centers in metal-coordinating groups—can thus be distinguished during the cross-modal interaction with RNA, through their separate atom representations.

Formally, the molecular graph is constructed as an attributed graph where atoms are nodes and bonds are edges, with atomic features (element type, charge, pharmacophore type, scaffold membership) and bond features (bond order, conjugation, stereochemistry) as node and edge attributes, respectively. A three-layer graph attention network computes a representation for each atom by aggregating information from its chemical neighborhood (Brody et al., 2022). The attention score between atoms *i* and *j* is:

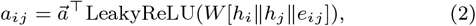

where *h*_*i*_ and *h*_*j*_ are the feature vectors of atoms *i* and *j, e*_*ij*_ is the edge feature vector, *W* is a learnable matrix, and ∥ denotes concatenation. The attention coefficient is normalized via:

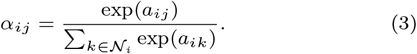

Three such layers are stacked to produce the final ligand embedding *H*_Ligand_ ∈ ℝ^*N*×*D*^, where *N* is the number of atoms and *D* is the hidden dimension.

### 2.5. Cross-Attention Interaction Module

As illustrated in Figure 1C, the cross-attention module enables RNA nucleotides to attend to ligand atoms through a 3-layer multi-head cross-attention mechanism (4 heads per layer) (Vaswani et al., 2017). In each layer, the RNA feature matrix *H*_RNA_ ∈ ℝ^*L*×*D*^ serves as queries, and the ligand atom matrix *H*_Ligand_ ∈ ℝ^*N*×*D*^ serves as keys and values, where *L, N*, and *D* denote the RNA sequence length, number of ligand atoms, and hidden dimension, respectively. For head *r*, linear projections produce *Q*_*r*_ = *H*_RNA_*W*_*Q,r*_, *K*_*r*_ = *H*_Ligand_*W*_*K,r*_, and *V*_*r*_ = *H*_Ligand_*W*_*V,r*_, with per-head dimension *d*_*h*_ = *D/*4. The scaled dot-product attention is:

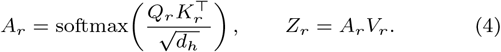

RNA residues aggregate ligand information through the concatenated head outputs, *H*_agg_ = Concat(*Z*_1_, …, *Z*_4_)*W*_*O*_. Ligand padding positions are masked before softmax. Each layer applies residual connections, layer normalization, and a feed-forward network:

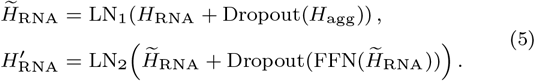

The feed-forward network has dimensions *D* → 4*D* → *D* with ReLU activation. Stacking three interaction layers progressively refines nucleotide representations based on ligand chemical properties (hydrogen-bond donors/acceptors, aromaticity, charge). These attention weights describe learned feature associations, rather than physical distances or docking trajectories.

### 2.6. Model Training and Optimization

#### 2.6.1. Training Objective and Probability Aggregation

As illustrated in Figure 1D, the final predictions are optimized through a probability-space weighted aggregation strategy. Binding site prediction is class-imbalanced: binding nucleotides account for approximately 25% of the training corpus and 27% of the combined test sets. Standard binary cross-entropy (BCE) does not explicitly down-weight easily classified examples. We adopt focal loss (Lin et al., 2017) to address this:

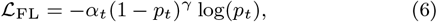

where *p*_*t*_ ∈ [0, 1] is the probability assigned to the true class, *γ >* 0 down-weights well-classified examples of either class, and *α*_*t*_ ∈ [0, 1] weights the two classes. For binding probability *p* and true label *y* ∈ {0, 1},

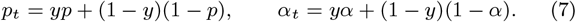

Empirically, *γ* = 2 and *α* = 0.25 in all experiments, and the loss is averaged over non-padding nucleotide positions. Building on this objective, two models (Model A and Model B) are trained independently with different random initializations. A grid search over the weight space (*ω*_*A*_, *ω*_*B*_) with *ω*_*A*_, *ω*_*B*_ ≥ 0 and *ω*_*A*_ + *ω*_*B*_ = 1 is performed on the validation set to identify the optimal weighting. The aggregated probability for each nucleotide is *p*_ens_ = *ω*_*A*_*p*_*A*_ + *ω*_*B*_*p*_*B*_, and the final prediction is determined by the threshold that maximizes F1 on the validation set.

### 2.7. Evaluation Metrics

To comprehensively evaluate binding site prediction performance, we adopt standard binary classification metrics: Precision (Prec), Recall (Rec), F1-score, and Matthews Correlation Coefficient (MCC) (Sokolova and Lapalme, 2009; Boughorbel et al., 2017; Chicco et al., 2021):

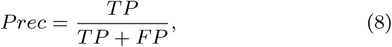

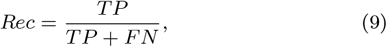

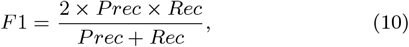

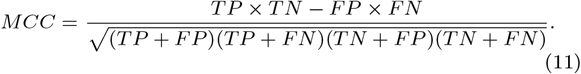

Here, TP and TN denote correctly identified binding and non-binding nucleotides, respectively, while FP and FN represent false alarms and missed binding sites. Precision quantifies the fraction of predicted binding sites that are genuine, and Recall measures the proportion of true binding sites successfully retrieved. The F1-score, as the harmonic mean of Precision and Recall, provides a balanced assessment. Given the class imbalance between binding and non-binding classes in RNA sequences, MCC is particularly informative because it yields a robust measure of classification quality even when class sizes are highly unequal. We also report the area under the receiver operating characteristic curve (AUROC, labeled AUC in the tables) and the area under the precision– recall curve (AUPRC) to evaluate performance across classification thresholds (Saito and Rehmsmeier, 2015).

## 3. Results

### 3.1. Comparison with Existing Methods on TE9 and TE18

We perform a systematic comparison of LigRNA against existing published methods for RNA-ligand binding site prediction on the TE9 and TE18 benchmark datasets. The baseline methods include geometry-based approaches (Rsite (Zeng et al., 2015) and Rsite2 (Zeng and Cui, 2016)), the network-based method RBind (Wang et al., 2018), the random forest method RNAsite (Su et al., 2021), and deep learning frameworks (RLBind (Wang et al., 2023) and GATRsite (Sun et al., 2025)).

The results on the TE9 dataset (Table 1) demonstrate that LigRNA outperforms all compared methods across every evaluation metric, achieving the highest Precision (0.80) among all compared methods (Fig. 3A).

**Table 1.** Performance comparison of LigRNA with existing methods on the TE9 dataset.

| Model | MCC | F1 | Prec | Recall | AUC |
| --- | --- | --- | --- | --- | --- |
| Rsite2 | 0.07 | 0.39 | 0.38 | 0.19 | 0.53 |
| Rsite | 0.16 | 0.25 | 0.43 | 0.35 | 0.58 |
| RBind | 0.28 | 0.35 | 0.68 | 0.23 | 0.59 |
| RNAsite | 0.43 | 0.50 | 0.75 | 0.38 | 0.79 |
| RLBind | 0.55 | 0.66 | 0.75 | 0.60 | 0.85 |
| GATRsite | 0.57 | 0.68 | 0.77 | 0.61 | 0.85 |
| **LigRNA** | **0.63** | **0.73** | **0.80** | **0.67** | **0.87** |

**Fig. 3:**
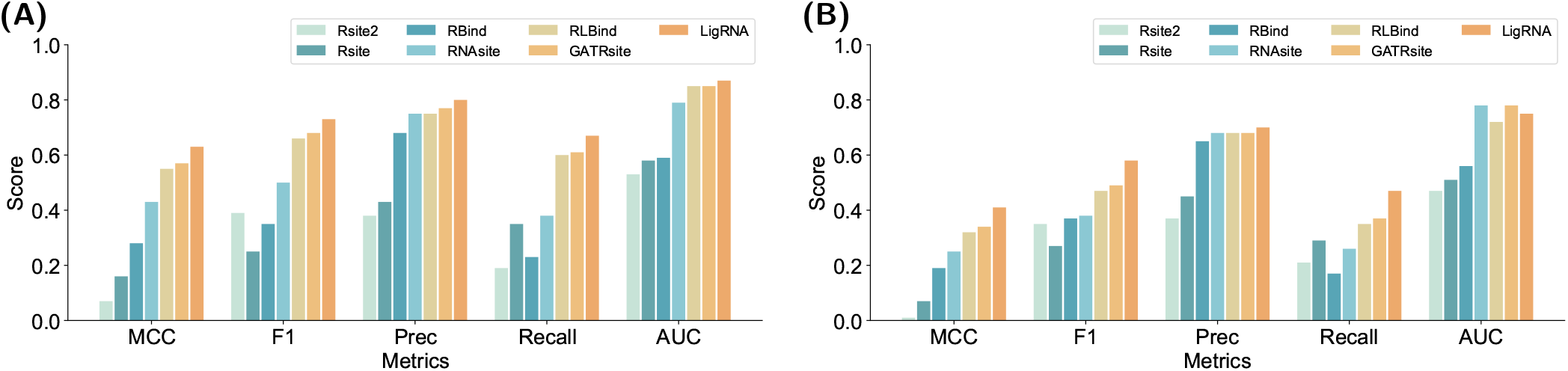
Performance comparison of all methods on TE9 (A) and TE18 (B) datasets. Bar charts show MCC, F1, Precision, Recall, and AUC for Rsite2, Rsite, RBind, RNAsite, RLBind, GATRsite, and LigRNA. LigRNA achieves the highest scores across most metrics on both datasets.

On the TE18 dataset (Table 2), most methods experience performance degradation relative to TE9, yet LigRNA maintains an advantage in F1 and MCC. After aggregation optimization, LigRNA achieves F1 = 0.58, MCC = 0.41, Precision = 0.70, Recall = 0.47 (Fig. 3B), outperforming GATRsite (F1 = 0.49, MCC = 0.34). However, its AUROC of 0.74 is below the 0.78 achieved by both GATRsite and RNAsite; the improvement does not extend to every metric.

**Table 2.** Performance comparison of LigRNA with existing methods on the TE18 dataset.

| Model | MCC | F1 | Prec | Recall | AUC |
| --- | --- | --- | --- | --- | --- |
| Rsite2 | 0.01 | 0.35 | 0.37 | 0.21 | 0.47 |
| Rsite | 0.07 | 0.27 | 0.45 | 0.29 | 0.51 |
| RBind | 0.19 | 0.37 | 0.65 | 0.17 | 0.56 |
| RNAsite | 0.25 | 0.38 | 0.68 | 0.26 | 0.78 |
| RLBind | 0.32 | 0.47 | 0.68 | 0.35 | 0.72 |
| GATRsite | 0.34 | 0.49 | 0.68 | 0.37 | 0.78 |
| **LigRNA** | **0.41** | **0.58** | **0.70** | **0.47** | 0.74 |

### 3.2. Ablation Study of Features and Architecture

To assess the contribution of each architectural component, we systematically ablated four key modules. Specifically, we examined the effects of replacing cross-attention with ligand mean pooling and concatenation (w/o. Attention), removing the RNA residual blocks while retaining the input projection (w/o. ResNet), removing explicit physicochemical features of ligand atoms (w/o. ChemEnv), and removing auxiliary RNA features including SASA and secondary structure (w/o. AuxFeatures). All models were trained with focal loss.

On the TE9 dataset, the full model achieved AUROC (0.87) and AUPRC (0.70). Inspection of the per-RNA F1 distribution (Fig. 4A) revealed distinct changes across variants. The most severe degradation occurred upon removal of the RNA ResNet module: the F1 distribution shifted markedly leftward (mean: 0.54 vs. 0.68 for the full model), indicating lower per-RNA performance on some complexes. Ablation of cross-attention (mean F1: 0.60) and auxiliary features (mean F1: 0.63) caused more moderate shifts, while removal of chemical environment features (mean F1: 0.62) resulted in an elevated proportion of mid-range F1 samples.

**Fig. 4:**
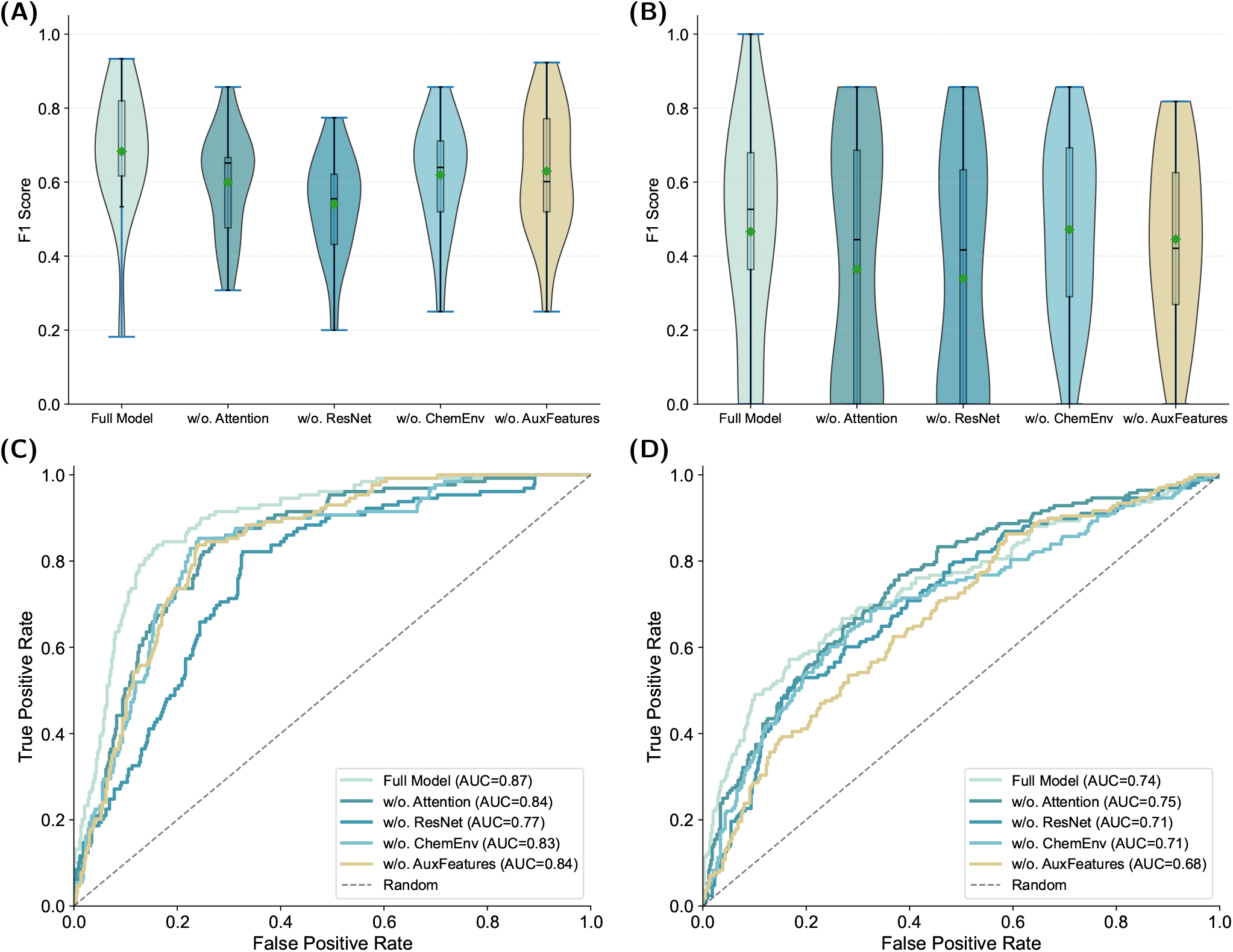
Per-RNA performance distribution of LigRNA. **(A, B)** Violin plots of per-RNA F1 scores on the TE9 (A) and TE18 (B) test sets, comparing the full model against four ablated variants. **(C, D)** Corresponding ROC curves for the full model and all four ablated variants on TE9 (C) and TE18 (D).

On the TE18 dataset, overall performance was weaker than on TE9, and per-RNA variance was considerably larger (Fig. 4B). Removal of the ResNet module drove F1 further down (mean: 0.34, std: 0.33), indicating substantial variability across RNAs. Notably, ablation of chemical environment features yielded F1 mean (0.47) comparable to or exceeding that of removing cross-attention (0.36) or auxiliary features (0.45), suggesting that the relative contributions of these features may vary across RNAs. Attributing these differences to metal-ion binding requires a separate analysis by ligand class.

To quantify statistical significance, we performed paired Wilcoxon signed-rank tests between the full model and its four ablated variants (Fig. 5A-D). On TE9, ablation of the RNA ResNet module resulted in the most pronounced degradation (*p*=0.0002 for F1, *p*=0.0006 for MCC, MCC reduction of 0.24), followed by removal of cross-attention and chemical environment features (*p*<0.01). The w/o. AuxFeatures variant showed no significant difference from the full model (*p*>0.05), providing no statistically significant evidence for a benefit from the auxiliary RNA features on this dataset. On TE18, only ablation of the RNA ResNet module reached statistical significance (*p*<0.05), while the remaining three variants did not exhibit significant differences, indicating that independent benefits from these components are not statistically established on TE18.

**Fig. 5:**
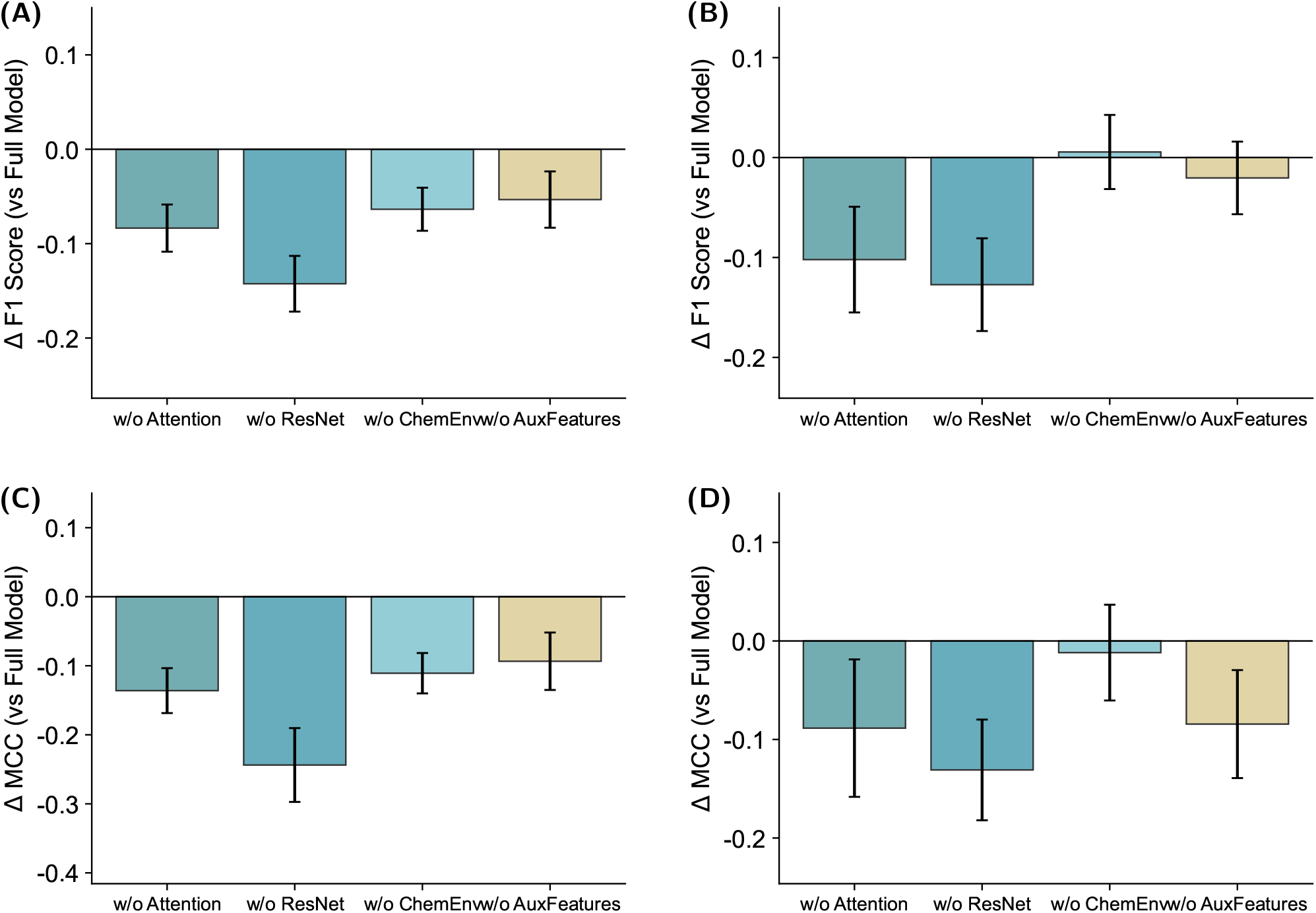
Ablation score differences for LigRNA. **(A, B)** ΔF1 between each ablated variant and the full model on TE9 (A) and TE18 (B), with statistical significance assessed by the Wilcoxon signed-rank test. **(C, D)** ΔMCC on TE9 (C) and TE18 (D). Differences are calculated as the ablated variant minus the full model; negative values indicate lower scores after ablation.

### 3.3. Case Studies

We examined two representative multi-ligand complexes from the benchmark datasets. The 2TOB structure (TE9) features a TPP riboswitch that simultaneously accommodates three ligands (2TB, TOA, and TOC) within a single RNA scaffold, with overlapping binding interfaces at nucleotides A:7–11, A:14–15, and A:17–18. Despite this multi-ligand complexity, LigRNA achieved perfect prediction accuracy for all three ligands in 2TOB (Fig. 6A, B). The 1PBR complex harbors three ligands (PA1, CYY, and IDG) occupying spatially distinct but adjacent sites. PA1 spans two binding zones (A:4–8 and A:17–24) encompassing 13 contact nucleotides, with partial overlap with CYY and IDG. LigRNA successfully recovered the complete PA1 binding interface (Fig. 6C, D). Together, these selected examples illustrate ligand-conditioned predictions on the same RNA scaffold, but do not isolate the contribution of cross-attention.

**Fig. 6:**
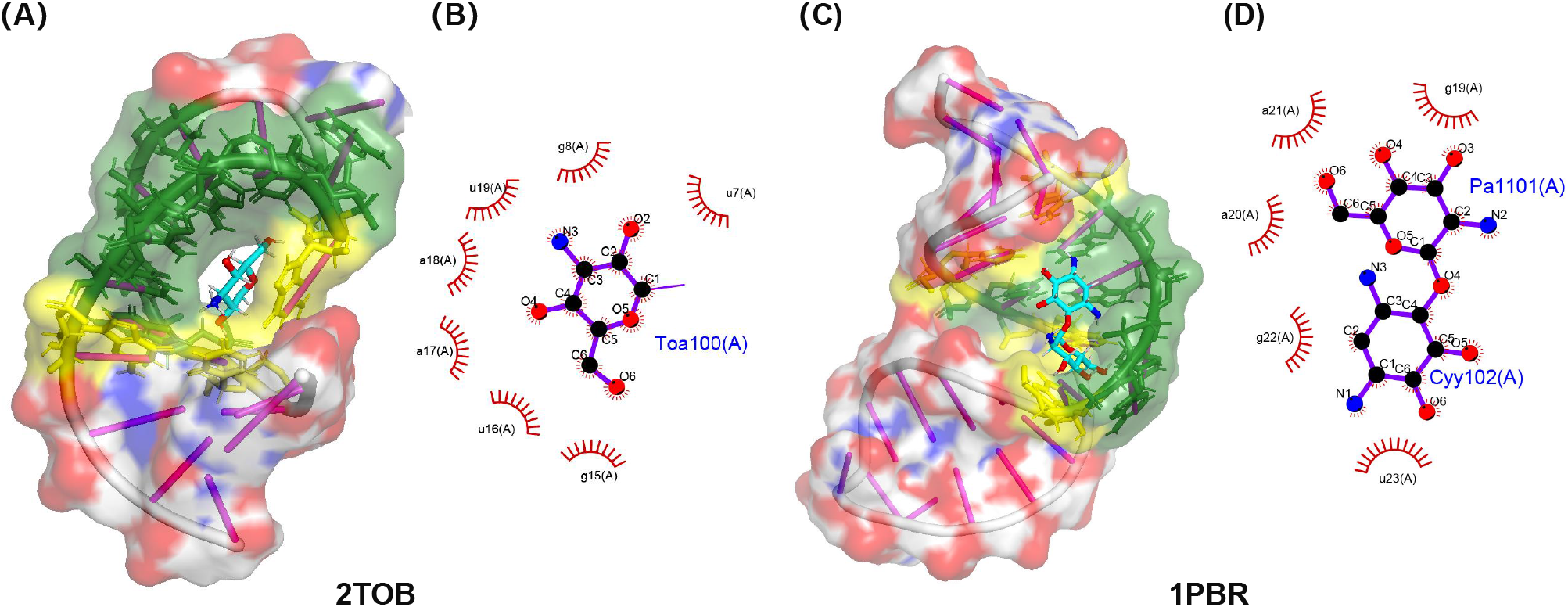
2D ligand-pocket interaction diagrams for the tobramycin-RNA aptamer complex (PDB ID: 2TOB) and the 16S ribosomal RNA A-site in complex with paromomycin (PDB ID: 1PBR). (A) 2TOB PyMOL: spatial arrangement of three co-bound ligands (2TB, TOA, TOC) within the same RNA cavity, with predicted binding sites shown in green and yellow. (B) 2TOB LigPlot: all seven binding residues (A:7-15) within 3 Å are correctly predicted. (C) 1PBR PyMOL: binding nucleotides of three co-bound ligands (PA1, CYY, IDG), with predicted binding sites shown in green and yellow. (D) 1PBR LigPlot: binding residues of the representative ligand PA1, spanning the 5’-proximal (A:4–8) and 3’-proximal (A:17–24) regions. Ligand names are colored in blue and ligand bonds in purple. The hydrophobic nucleotides of pockets and hydrophobic interactions are colored in red.

To evaluate the model’s capacity to generalize to RNA families absent from both training and test sets, we analyzed the Pepper RNA aptamer complexed with the fluorogenic ligand HBC514 (PDB ID: 7EON). The Pepper aptamer adopts a tuning-fork-like topology (Huang et al., 2021) distinct from those in TE9 or TE18, and HBC514 represents a chemically novel ligand chemotype absent from all training datasets, providing an illustrative generalization case. At a 4.0 Å threshold, LigRNA correctly predicted all binding nucleotides in 7EON with zero false positives and zero false negatives (Fig. 7), showing accurate prediction for this complex. A single case does not establish generalization across RNA families or ligand chemotypes, or identify the physical interactions learned by the model.

**Fig. 7:**
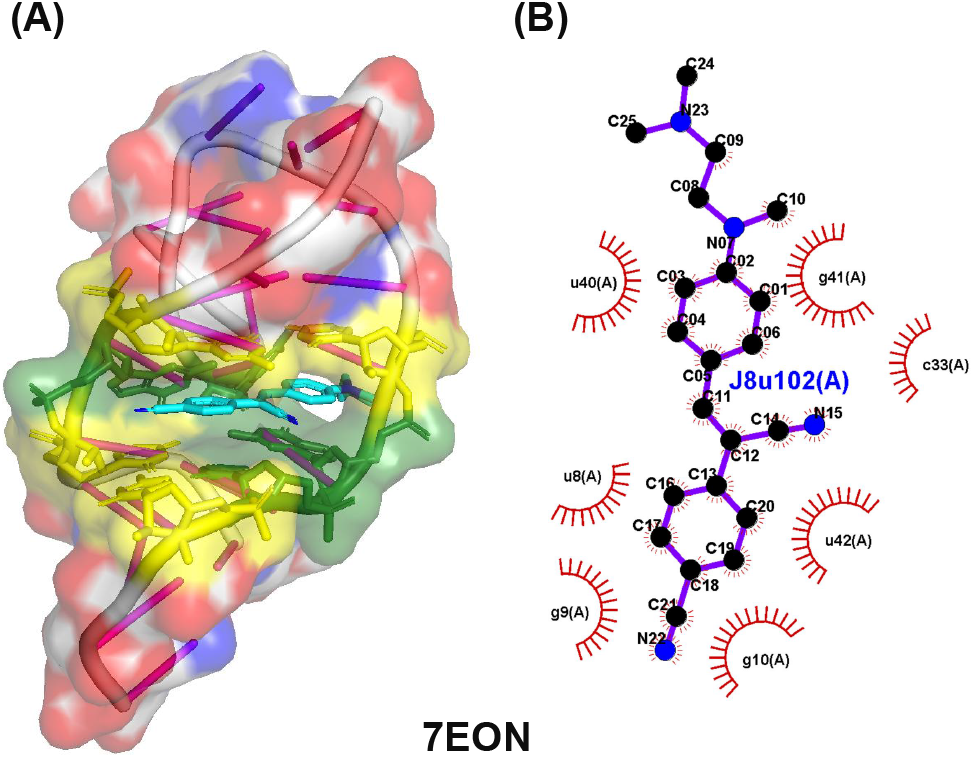
Generalization analysis on the 7EON Pepper aptamer. (A) PyMOL-rendered structure with HBC514 ligand in the fork-like aptamer pocket; binding nucleotides in green. (B) LigPlot diagram showing the hydrogen-bonding network and *π*-*π* stacking interactions between HBC514 and the aptamer. LigRNA achieves correct prediction on all binding nucleotides at a 4.0 Å threshold on this out-of-sample complex.

## 4. Discussion

On the TE9 dataset, LigRNA achieves an F1 of 0.73, surpassing GATRsite (F1 = 0.68) by 7.4%. The most notable gain is in Recall, which increases from 0.61 to 0.67, indicating that LigRNA retrieves a substantially larger fraction of native binding nucleotides without proportionally increasing false positives. Two complementary factors may contribute to this gain. First, the cross-attention mechanism enables nucleotides to attend selectively to chemically relevant ligand atoms, allowing ligand-conditioned nucleotide representations. Second, the SASA and secondary structure features provide information about nucleotide exposure and local topology that may help control false positives. However, the auxiliary-feature ablation does not establish an independent benefit on TE9.

The TE18 results reveal a different challenge profile. Most methods show lower scores on TE18, with LigRNA showing a drop in Recall from 0.67 to 0.47. Differences in RNA families and ligand classes, including metal ion-RNA complexes, may contribute to this pattern. Metal-ion binding involves coordination chemistry that differs from many organic small-molecule interactions. Our secondary structure analysis (Fig. 2) describes the TE9 binding sites: knotted structures and internal loops together account for 55.4% of binding site nucleotides. These percentages alone cannot explain performance differences between TE9 and TE18. Whether metal-ion sites differ in these structural categories requires a separate analysis. The observation that ablation of chemical environment features yields comparable or higher F1 than ablation of cross-attention does not by itself establish a difference between ligand classes. These observations motivate further investigation of dedicated encoding strategies for metal coordination chemistry, such as incorporating ionic radius, coordination geometry, and ligand field stabilization energy.

The three-part case study further contextualizes these quantitative results. The 2TOB and 1PBR examples from TE9 illustrate that the model maintains high prediction accuracy even when multiple chemically distinct ligands occupy overlapping or spatially adjacent binding pockets on the same RNA scaffold – a scenario that reflects the complexity of real RNA-ligand interaction landscapes. The 7EON Pepper aptamer analysis provides an example involving an RNA family absent from both training and test sets: the correct prediction on all binding nucleotides at a 4.0 Å threshold shows accurate prediction for this complex, although one case cannot establish broader generalization. Such predictive performance, if confirmed across broader datasets, could be useful for fragment-based drug discovery and RNA-targeted therapeutic optimization on emerging targets, where high-precision binding-site predictions may help prioritize downstream experimental validation.

Despite these demonstrated improvements, several limitations should be considered. First, performance variance across individual RNAs remains large on TE18 (F1 std = 0.30), indicating that LigRNA excels on some complex families while struggling on others. This per-RNA heterogeneity may reflect the inherent diversity of RNA-ligand binding modes and underscores the need for more comprehensive benchmark datasets. Second, all benchmarks are derived from experimentally determined structures, which may introduce a bias toward experimentally tractable RNA-ligand complexes and away from more flexible or transient binding events. Third, the current model does not distinguish between binding site nucleotides that make energetically important contacts and those that are merely spatially close to the ligand. Incorporating explicit interaction-type annotations in future training data could enable more nuanced predictions that better support medicinal chemistry efforts.

In summary, the design principles examined here may be relevant beyond LigRNA itself: further advances in binding site prediction may benefit from simultaneous progress on both representation and interaction modeling. Richer ligand encodings, structure-aware RNA features, and dynamic cross-modal attention provide a framework for learning associations between RNA and ligand features, and these principles may inform the design of future architectures in this domain.

## 5. Conclusion

We presented LigRNA, a graph neural network framework for high-precision prediction of RNA-ligand binding sites from RNA sequences and small-molecule structures. By integrating deep evolutionary semantics from the structRFM foundation model with explicit secondary structure elements and solvent accessibility features, LigRNA constructs a comprehensive representation of the RNA local microenvironment. On the ligand side, the adoption of attributed molecular graphs and the GATv2 mechanism enables the model to capture atom-level chemical environments and pharmacophore attributes. The 3-layer multi-head cross-attention mechanism replaces static feature fusion, enabling the model to learn input-dependent associations between RNA residues and ligand functional groups. Evaluations on the TE9 and TE18 benchmark datasets show higher F1 and MCC than the compared methods, although TE18 AUROC is below that of GATRsite and RNAsite. Ablation studies identify the RNA residual encoder as the most consistent contributor, while evidence for other components varies by dataset. LigRNA provides a computational tool for identifying RNA-ligand interaction interfaces, with potential to accelerate the discovery and optimization of RNA-targeted therapeutics.

## 6. Data availability

Dataset composition and PDB identifiers are provided in Supplementary Tables S1 and S2. The corresponding source structures can be obtained from the Protein Data Bank at https://www.rcsb.org/.

## 7. Code availability

The source code used in this study is freely available at https://github.com/WillowWang0216/LigRNA.

## 8. Supplementary data

Supplementary material accompanies this article.

## 9. Author contributions statement

P.X. and S.K.Z. conceptualized and administered the study. H.Z. co-supervised the study and designed the framework and experiments. X.W. implemented the framework and conducted the experiments. X.W. and H.Z. analyzed the results and wrote the paper. H.L. and A.C. analyzed the comparative experimental results. F.Z. participated in the design and parameter tuning of aggregation strategies. H.C. and C.W. collected and processed the datasets. K.C. arranged computational resources. All authors read, contributed to the discussion, and approved the final paper.

