## Supplementary for "LigRNA: High-accuracy prediction of RNA-ligand binding sites via dynamic chemical complementarity and language model encoding"

*Bioinformatics*, 2026, 1–6
Supplementary Material

### Supplementary Material for LigRNA: High-accuracy prediction of RNA-ligand binding sites via dynamic chemical complementarity and language model encoding

**Xin Wang,[1,2,†] Heqin Zhu,[1,2,†,*] Hao Li,[1,2] Ao Chang,[1,2] Feng Zhang,[1,2] Haobin Chen,[1,2] Chenxi Wang,[1,2] Ke Chen,[1,2] Shaohua Kevin Zhou[1,2,3,4,5,*] and Peng Xiong[1,2,4,*]**

[1]School of Biomedical Engineering, Division of Life Sciences and Medicine, University of Science and Technology of China (USTC), Anhui, 230026, China, [2]Medical Imaging, Robotics, Analytic Computing & Learning (MIRACLE) Lab, YRD-RIGHT, Suzhou Institute for Advance Research, USTC, Jiangsu, 215123, China, [3]Jiangsu Key Laboratory of Multimodal Digital Twin Technology, Jiangsu, 215123, China, [4]Biomedical Basic Research Center (BBRC) of Jiangsu, Jiangsu, 215123, China and [5]State Key Laboratory of Precision and Intelligent Chemistry, Anhui, 230026, China

##### Contents

Dataset composition and PDB identifiers (Tables S1 and S2); ligand atom and bond features (Tables S3 and S4); ResNet depth analysis (Tables S5 and S6); model-level metric heatmaps for BCE and focal loss (Fig. S1); per-RNA F1 distributions with BCE loss (Fig. S2); and paired ablation comparisons (Fig. S3).

© The Author 2026. Published by Oxford University Press. All rights reserved. For permissions, please 

#### 1. Supplementary Tables

##### 1.1. Dataset Construction

The benchmark datasets used for training and evaluation are derived from public repositories of RNA-ligand complex structures. RNA-ligand complexes were retrieved from the PDB as of October 2025. Complexes were retained if they contained at least one RNA chain and one small-molecule ligand with molecular weight $\leq$ 1000 Da; crystalline water molecules were excluded. Binding-site annotations follow the benchmark preprocessing protocols described in the main text. For the structural case illustrations, the contact cutoffs are 3 Å for 2TOB and 4.0 Å for 7EON, as specified in the corresponding main-text figure captions. These case-specific contact cutoffs are distinct from the probability threshold used to classify model predictions. RNA chains were simplified by manually inspecting PDB structures to remove structurally redundant segments, ensuring that each complex contains the minimal necessary set of RNA chains. Structural redundancy among RNA chains was eliminated using TM-align with a TM-score threshold of 0.3; chains with TM-score $\geq$ 0.3 to an existing cluster representative were assigned to that cluster, otherwise initiating a new cluster. To enrich the training corpus, complexes with TM-score below 0.5 relative to existing clusters were retained, thereby increasing the diversity of micro-chemical interaction environments in the training set. Cluster overlap between Train60, TE18, and TE9 was eliminated by enforcing a TM-score threshold of 0.3 across all three sets, preventing data leakage. This procedure yielded 228 additional PDB structures (Train228, 241 chains, 67 clusters), which were merged with Train60 and split 8:2 into training and validation partitions. Full dataset composition and binding-site statistics are provided in Supplementary Table S1, and the complete list of PDB structure identifiers is provided in Supplementary Table S2. Notably, the training corpus spans a broad spectrum of RNA-ligand interaction types: the bound small molecules encompass over 150 distinct ligand chemotypes, including metal ions (e.g., $Mn^{2+}$, $Mg^{2+}$), nucleotides and nucleotide analogs, aminoglycosides, macrolides, cofactor derivatives, polyamines, and fluorogenic probes.

**Table S1.** Summary of RNA-ligand datasets used in this study. The total row is the arithmetic sum of the four dataset rows, without deduplication across datasets.

| Dataset | PDB Count | RNA Chain Count | Positive Samples | Negative Samples | Total Nucleotides |
|---|---|---|---|---|---|
| Train228 | 228 | 241 | 1,459 | 3,331 | 4,790 |
| Train60 | 60 | 75 | 467 | 2,372 | 2,839 |
| TE18 | 18 | 27 | 197 | 458 | 655 |
| TE9 | 9 | 9 | 131 | 434 | 565 |
| Total | 315 | 352 | 2,254 | 6,595 | 8,849 |

**Table S2.** Dataset composition and contained PDB structure IDs.

| Dataset | Contained PDB Structure IDs |
|---|---|
| Train228 | 1CX5,1F27,1FJG,1HNX,1I97,1J7T,1K73,1KC8,1KOD,1KQS,1LC4,1MWL,1N8R,1NJI,1NJM,1O9M,1Q7Y,1SM1,1VQ8,1VQN,1VQP,1XBP,1YIT,1YJN,1YJW,1ZZ5,2DQO,2E9R,2ESI,2ET3,2ET5,2F4U,2F4V,2FCX,2FCZ,2HHH,2HO6,2HOJ,2HOP,2KD4,2KMJ,2KU0,2L94,2N0J,2NZ4,2O3V,2O3W,2O3X,2OE8,2OGM,2OGN,2OTJ,2OTL,2PPB,2QEX,3B0V,3CXC,3DIR,3DVV,3EGZ,3F4G,3FWO,3G6E,3G71,3G96,3I55,3J7A,3JQ4,3L3C,3SUH,3TD1,3WRU,4AQY,4DR1,4DR2,4DV1,4E8M,4FAR,4FAU,4GPX,4II9,4KVB,4LF4,4LFA,4LFB,4LFC,4LVV,4LVY,4NYA,4P20,4P95,4PJO,4Q5V,4QVI,4WCE,4WFA,4WFB,4X4V,4YCO,4YCP,4Z0C,5AJ3,5BR8,5BS3,5BWS,5DM7,5HL7,5IWA,5KPY,5ME0,5NRG,5NS3,5NS4,5V1K,5V7Q,5WNT,5X21,5XI1,5Z1H,5Z71,5ZC9,5ZEJ,6AZ1,6BFB,6BJY,6C63,6C64,6CAQ,6D2U,6DB8,6DDD,6DDG,6DN2,6E1S,6E1T,6E1U,6E1V,6E82,6E84,6E8S,6E8T,6G2K,6GZK,6HC5,6HMO,6IZP,6JBG,6JJH,6PC5,6PC6,6PCH,6PCQ,6PCR,6PQ7,6QIT,6QIV,6QUL,6TFF,6UC9,6UP0,6V9B,6V9D,6VA2,6VA3,6VA4,6WQN,6WQQ,6WRS,6WRU,6WZR,6WZS,6XB7,6XJQ,6XJW,6XKI,6YSI,6ZV6,7A18,7A3Y,7AAV,7ABI,7BPF,7D12,7D7V,7D81,7DUG,7DUH,7DUI,7DUJ,7E9E,7FJ0,7KKV,7KVT,7KVV,7L0Z,7MKT,7OA3,7OAW,7OOD,7RWR,7TZR,7TZS,7TZT,7U0Y,7WIF,7WII,8A5I,8ANA,8AS7,8CA7,8CAI,8CEU,8CGD,8CGJ,8CGU,8CGV,8CVM,8D2A,8D2B,8D5L,8D5O,8EYU,8EYV,8EYW,8GXB,8HB3,8HBA,8R62 |
| Train60 | 5J02,5O69,5U3G,5VCI,6FZ0,6HAG,1NTB,1QD3,1RAW,1TOB,1UTS,1UUD,1Y90,1YLS,1YRJ,2FCY,2G5K,2KGP,2KTZ,2L8H,2LWK,2MXS,2NOK,2QUW,2YIE,3BNQ,3C44,3FU2,3GX3,3MEI,3OXE,3Q3Z,3Q50,3SKT,3TZR,3VRS,4FRG,4K31,4LVX,4MEG,4MGM,4NYB,4PCJ,4QJH,4QLM,4RGE,4XW7,5D5L,5DH8,5FJ1,1AJU,1ARJ,1BYJ,1EHT,1EI2,1F1T,1FYP,1HR2,1LVJ,1NBK |
| TE18 | 1DDY,1F1T,1FMN,1NEM,1Q8N,2JUK,2MIS,2PWT,2TOB,364D,379D,430D,4F8U,4PQV,4YAZ,5BJO,5V3F |
| TE9 | 1NEM,1PBR,1Q8N,1UUI,1Y26,2GDI,2TOB,3D2X,3GX2 |

**Table S3.** Atomic features used in the ligand model.

| Atomic Feature | C0 | N4 | O5 | Description |
|---|---|---|---|---|
| atomic_number | 1 | 2 | 3 | atomic number |
| chirality | 0 | 0 | 0 | chirality |
| degree | 3 | 3 | 1 | degree |
| formal_charge | 0 | 0 | 0 | formal charge |
| num_H | 0 | 2 | 0 | number of hydrogen atoms |
| num_radical_e | 0 | 0 | 0 | number of radical electrons |
| hybridization | 2 | 2 | 1 | hybridization state |
| is_aromatic | 0 | 0 | 0 | aromaticity |
| is_in_ring | 0 | 0 | 0 | ring membership |
| Hbond_donor | 0 | 1 | 0 | hydrogen bond donor |
| Hbond_acceptor | 0 | 1 | 1 | hydrogen bond acceptor |
| Basic | 0 | 1 | 0 | basic character |
| Acid | 1 | 0 | 0 | acidic character |
| Halogen | 0 | 0 | 0 | halogen atom |
| Scaffold | 0 | 0 | 0 | Murcko scaffold membership |

**Table S4.** Edge (bond) features used in the ligand model.

| Bond Feature | Value | Description |
|---|---|---|
| bond_type | 0=single, 1=double, 2=triple, 3=aromatic | chemical bond type |
| bond_stereo | 0=none, 1=Z (cis), 2=E (trans) | stereochemistry configuration |
| is_conjugated | 0=no, 1=yes | whether the bond is in a conjugated system |
| atom_pair | 0=C–C, 1=C–N, 3=C–O, . . . , 14=other | element pair of bonded atoms |

##### 1.2. Analysis of ResNet Depth

To investigate the impact of the number of residual blocks in the RNA sequence encoder on prediction performance and identify the optimal architecture configuration, we trained models with the ResNet depth parameter (*rna_depth*) ranging from 1 to 9, corresponding to architectures denoted as ResNet-1 through ResNet-9 (each integer denotes the number of stacked residual blocks, where each block contains two convolutional layers). A fixed sigmoid threshold of 0.5 was applied during inference. As summarized in Table S5 (TE9) and Table S6 (TE18), performance on TE9 generally improves as depth increases from 1 to 5 residual blocks, with ResNet-5 achieving the best overall results (MCC=0.51, F1=0.63, AUC=0.84). Beyond 5 blocks, performance fluctuates and declines: ResNet-7 shows the most pronounced degradation (F1=0.51, AUC=0.73), showing that greater depth does not consistently improve performance on TE9. On TE18, performance is more volatile across depths without a clear monotonic trend. ResNet-5 achieves the highest F1 (0.48), whereas ResNet-9 achieves the highest AUC (0.72). Deeper architectures such as ResNet-7 and ResNet-9 maintain reasonable Precision but exhibit substantially lower Recall, leading to overall metric deterioration. This indicates that greater depth does not consistently improve binding-site coverage on TE18. Based on these observations, ResNet-5 (five residual blocks) was adopted as the default backbone configuration in all single-model experiments, including the ablation study reported in main-text Section **??**. The final ensemble model used for comparison with baseline methods also adopts ResNet-5 as its backbone.

**Table S5.** Performance comparison of models with different ResNet depths (number of residual blocks) on the TE9 dataset.

| Model | MCC | F1 | Prec | Recall | AUC |
|---|---|---|---|---|---|
| ResNet-1 | 0.32 | 0.50 | 0.41 | 0.64 | 0.75 |
| ResNet-2 | 0.42 | 0.56 | 0.53 | 0.60 | 0.81 |
| ResNet-3 | 0.47 | 0.60 | 0.50 | 0.75 | 0.82 |
| ResNet-4 | 0.45 | 0.59 | 0.48 | 0.76 | 0.82 |
| ResNet-5 | 0.51 | 0.63 | 0.52 | 0.78 | 0.84 |
| ResNet-6 | 0.45 | 0.59 | 0.50 | 0.71 | 0.82 |
| ResNet-7 | 0.34 | 0.51 | 0.44 | 0.60 | 0.73 |
| ResNet-8 | 0.42 | 0.55 | 0.55 | 0.55 | 0.82 |
| ResNet-9 | 0.43 | 0.58 | 0.48 | 0.73 | 0.82 |

**Table S6.** Performance comparison of models with different ResNet depths (number of residual blocks) on the TE18 dataset.

| Model | MCC | F1 | Prec | Recall | AUC |
|---|---|---|---|---|---|
| ResNet-1 | 0.30 | 0.45 | 0.56 | 0.37 | 0.65 |
| ResNet-2 | 0.24 | 0.45 | 0.46 | 0.43 | 0.69 |
| ResNet-3 | 0.23 | 0.40 | 0.49 | 0.34 | 0.67 |
| ResNet-4 | 0.27 | 0.43 | 0.53 | 0.36 | 0.68 |
| ResNet-5 | 0.29 | 0.48 | 0.50 | 0.45 | 0.71 |
| ResNet-6 | 0.30 | 0.45 | 0.56 | 0.38 | 0.70 |
| ResNet-7 | 0.24 | 0.38 | 0.53 | 0.30 | 0.71 |
| ResNet-8 | 0.29 | 0.44 | 0.54 | 0.38 | 0.66 |
| ResNet-9 | 0.20 | 0.30 | 0.53 | 0.21 | 0.72 |

#### 2. Supplementary Figures

##### 2.1. Comparison of BCE and Focal Loss

Figure S1 presents model-level metric heatmaps for the full model and four ablated variants trained with binary cross-entropy (BCE) or focal loss. Rows correspond to model variants and columns to evaluation metrics. These plots summarize model-level scores, rather than scores for individual RNA complexes. The per-RNA F1 distributions for BCE are shown separately in Fig. S2; the corresponding focal-loss distributions appear in main-text Fig. **??**A–B.

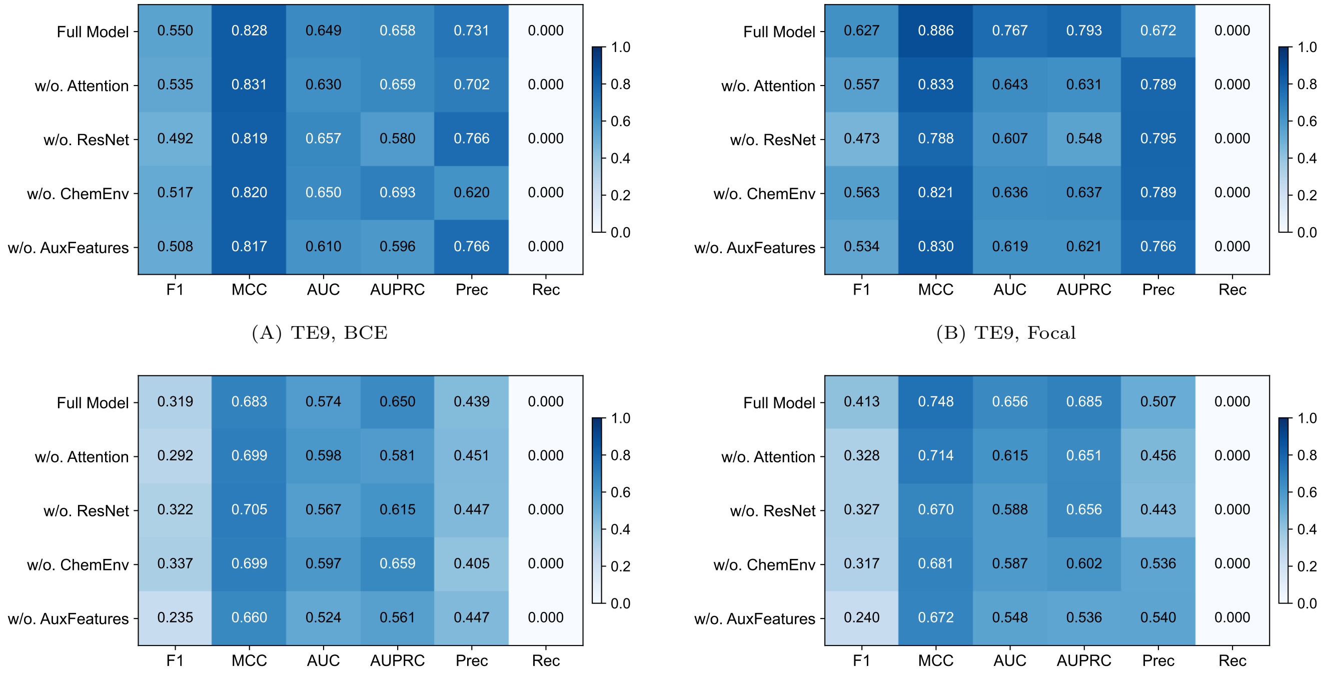


(A) TE9, BCE (B) TE9, Focal

(C) TE18, BCE (D) TE18, Focal

Fig. S1: **Model-level metric heatmaps for BCE and focal loss.** Panels show TE9 with BCE (A) or focal loss (B), and TE18 with BCE (C) or focal loss (D). Rows correspond to the full model and four ablated variants; columns are labeled F1, MCC, AUC, AUPRC, precision (Prec), and recall (Rec). Cell colors and annotations show the displayed metric values.

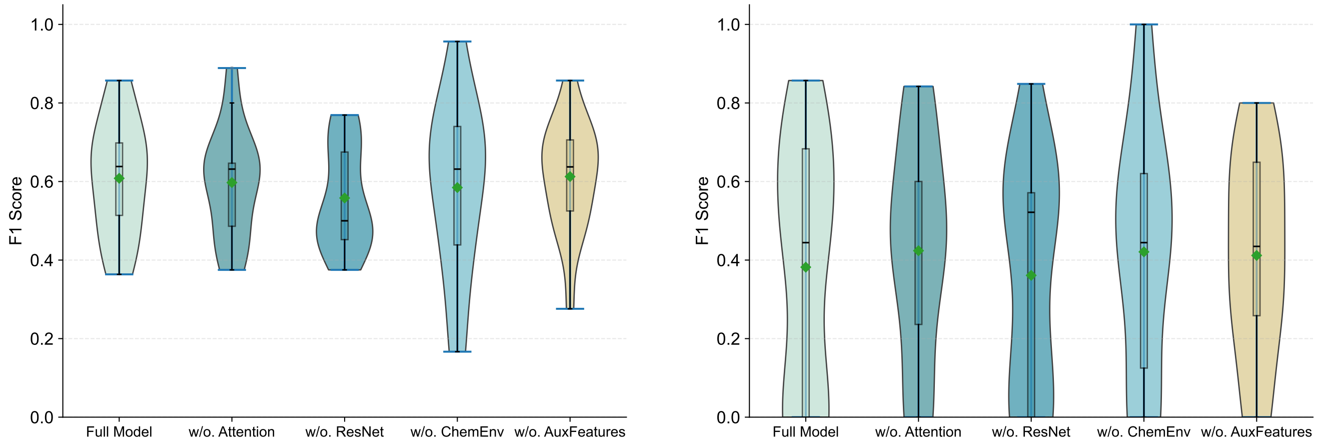


(A) TE9, BCE (B) TE18, BCE

Fig. S2: **Per-RNA F1 score distribution of models trained with BCE loss on TE9 and TE18 datasets.** Violin plots comparing the full model against four ablated variants trained with binary cross-entropy loss. See main-text Fig. **??**A–B for the corresponding results with focal loss.

#### 2.2. Ablation Significance Scatter Plots

To assess the statistical significance of the ablation results, we performed pairwise Wilcoxon signed-rank tests between the full model and each ablated variant on both datasets. The scatter plots directly compare per-RNA F1 and MCC scores of the full model against each ablated variant, with points above the diagonal indicating variants that outperform the full model (Fig. S3). On TE9, most ablated variants fall substantially below the diagonal for both F1 and MCC, illustrating lower scores for many paired comparisons. Ablation of the RNA ResNet module shows the most pronounced downward shift, consistent with its contribution to prediction performance. On TE18, the scatter distributions exhibit greater dispersion, reflecting the larger per-RNA performance variance on this more challenging dataset. Nevertheless, the ResNet ablation still consistently underperforms the full model across most samples, while the other three variants show more variable behavior, consistent with the Wilcoxon test results ($p$<0.05 for ResNet, $p$ >0.05 for the remaining variants).

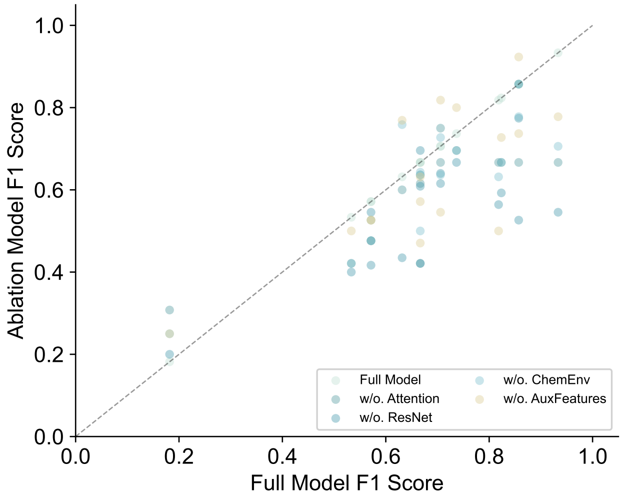


(A) TE9, F1

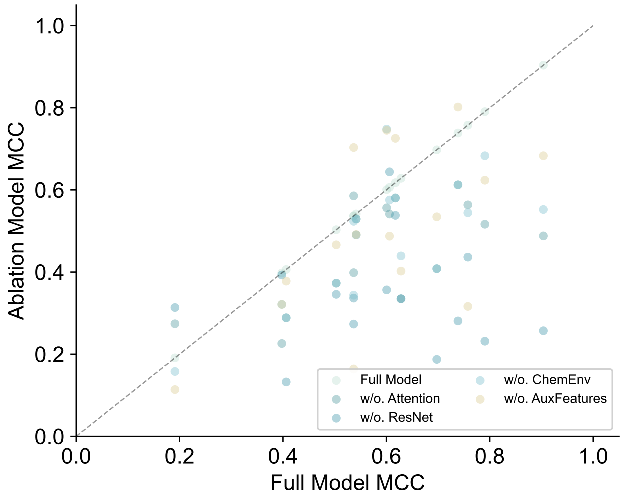


(B) TE9, MCC

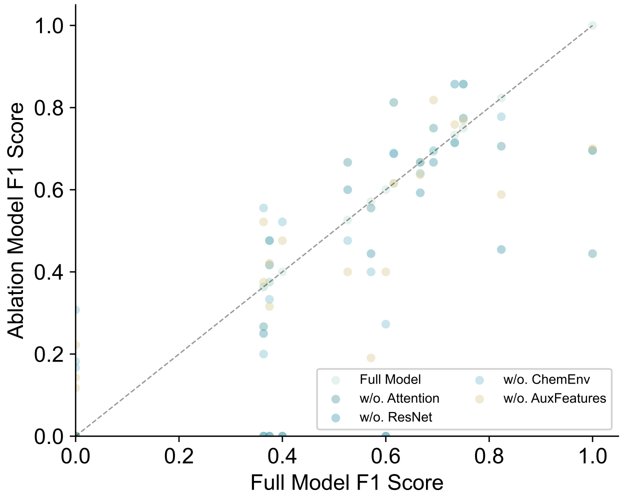


(C) TE18, F1

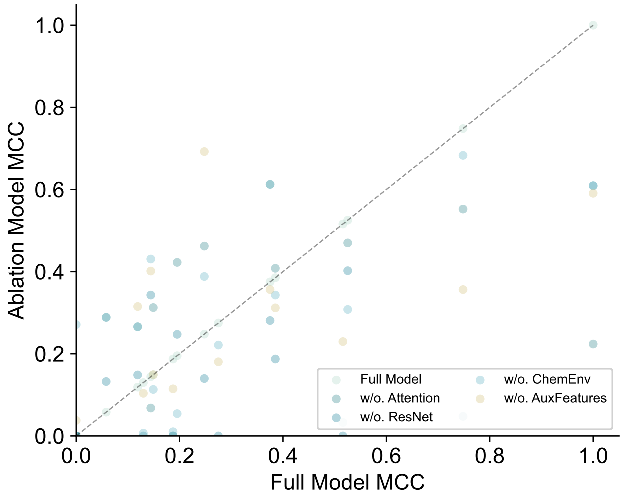


(D) TE18, MCC

Fig. S3: **Paired per-RNA scores for ablated variants and the full model.** Panels show F1 (A, C) and MCC (B, D) on TE9 (A, B) and TE18 (C, D). The horizontal axis gives the full-model score and the vertical axis the ablated-variant score. Points below the diagonal indicate lower scores for the ablated variant.
